# Biological responses to terahertz radiation with different power density in primary hippocampal neurons

**DOI:** 10.1101/2022.04.04.487008

**Authors:** Li Zhao, Ruhan Yi, Sun Liu, Yunliang Chi, Shengzhi Tan, Ji Dong, Hui Wang, Jing Zhang, Haoyu Wang, Xinping Xu, Binwei Yao, Bo Wang, Ruiyun Peng

## Abstract

**Background:** Terahertz (THz) radiation is a valuable imaging and sensing tool which is widely used in industry and medicine. However, it biological effects including genotoxicity and cytotoxicity are not clear, particularly on the nervous system. In this study, we investigated how THz radiation with different frequency and intensity would affect the morphology, cell growth and function of rat hippocampal neurons *in vitro*.

**Methods:** Primary hippocampal neuronal cells were isolated from newborn Wistar rat pups and were exposed to THz radiation with the frequencies of 0.12 THz and 0.141 THz, at an power intensities of 10 mW/cm^2^ and 30 mW/cm^2^ respectively. The cellular activities were evaluated by Cell Counting Kit-8 after exposure for 10 min and 30 min. The cellular apoptosis was examined by Annexin V staining using flow cytometry and the ultrastructure was detected by electron microscopy. Moreover, the amount of amino acid in the cultured neurons was measured by high performance liquid chromatography (HPLC).

**Results:** No obvious morphological changes, such as axon length and axon number, could be detected in primary neurons after exposure to 10mW/cm^2^ or 50 mW/cm^2^ THz for 30 min. However, the overall cellular activities were differentially affected by 10mW/cm^2^ and 50 mW/cm^2^ THz. Obvious cellular apoptosis was induced by both 10mW/cm^2^ and 50 mW/cm^2^ exposure. We also found that the amounts of most amino acids did not change significantly after exposure to these two types of THz radiation.

**Conclusions:** The effects of THz radiation on hippocampal neurons depend on frequency, power, exposure time as well as the measurement time after the exposure. These results will provide important information for further investigation of the effect of THz radiation on the nervous system.

## 1. Background

Terahertz (THz) waves are non-ionizing radiation with frequencies ranging from 0.1 to 10 THz and have wavelengths between 3 mm and 30 μm. THz waves have been emerged as valuable tools in various fields, such as biomedical imaging, industrial quality control, defense and security[1], due to its unique properties. Firstly, THz waves can penetrate through a wide variety of materials, such as skin, fabric, plastic and wood. Secondly, many particles including biomolecules such as proteins and DNA have characteristic absorption lines at frequencies within THz range. Thirdly, THz radiation is highly reflected by metals. Although THz radiation does not generate free radicals, the anxiety and speculation about potential health hazards risks and biological effects has been growing fast in recent years.

Recent years, more and more groups are interested in exploring the biological effects of THz radiation[2, 3]. Most of the studies focused on fibroblasts and lymphocytes. And, contradictory results are always been reposted, due to various parameters, such as frequency and power intensity. However, the THz radiation on neuronal cells and nervous system are rarely investigated. Bourne *et al* reported that 0.14THz radiation with intensity between 24 and 62 mW/cm^2^ does not produce detectable adverse effects in cell line [4]. Another study suggested that low power THz radiation does not cause detectable effect in primary neurons, while high power radiation changes membrane potential [5]. *In vivo* study showed that THz radiation increased level of depression, indicating that THz reduced neuronal activities [6].

In this study, we aimed to investigate THz caused biological effects in primary rat hippocampal neurons. We exposed primary neuron to THz wave, and the potential toxicity and apoptosis was examined by CCK-8 assay and Annexin V staining, respectively. We found that exposure to THz with two different power intensities differentially affected cellular activity and apoptosis. The effects were confirmed by examining ultrastructure of the cells. THz radiation also slightly altered the concentrations of certain amino acid neurotransmitters in the culture medium. However, THz radiation did not produce significant effects on neuron growth and differentiation. Our results indicated that the effects of THz radiation on hippocampal neuron depended on parameters of THz radiation, such as frequency, power, exposure time as well as the measurement time.

## 2. Methods

### 2.1 Primary rat hippocampal neurons

Primary hippocampal neuronal cells were isolated according to the published protocols [7]. Briefly, newborn Wistar rat pups were euthanized within 12hr after birth and sterilized briefly with 75% ethanol. Cranium of the pup was opened using sterile scissors and the entire brain was removed. The brain was rinsed with cold tissue dissociation buffer (HEPES with 0.3% glucose, 0.75% sucrose, 320-330mOsm, pH 7.2-7.4). Meninges surrounding the hippocampus was removed with forceps. The isolated hippocampus was gently minced in a tissue culture dish. The minced tissue was then transferred to HBSS buffer containing 0.25% Trypsin and incubated at 37°C for 15 to 20 min with gentle shaking every 5 min. Digested tissue was centrifuged for 5 min at 150g. The supernatant was removed and cells were counted using a hemacytometer. The experiments and corresponding procedures were approved by the ethics committee from Beijing Institute of Radiation Medicine.

Primary rat hippocampal neurons were cultured in DMEM medium with 10% FBS, 10% horse serum and 1% L-Glutamine at a density of 5×10^5^/ml. Four hours later, the medium was changed to DMEM plus 10% horse serum, 1% N-2, 2% B-27, 1% L-Glutamine. Ara-C (5 μg/ml) was added to the medium after 24 hours’ culture, and 50% of the total medium was changed every 24h from then on.

### 2.2 THz radiation

*In vitro* exposure was performed using THz radiation system from Beijing Institute of Radiation Medicine. The frequencies and power intensities were 0.12 THz/10 mW/cm^2^, 0.141 THz/30 mW/cm^2^ and 0.157 THz/50 mW/cm^2^. The radiation periods were 10 min and 30 min respectively. Cells in sham-irradiated group were processed as that in THz exposed group, but without radiation.

### 2.3 Cellular activity assay

The cellular activity was analyzed by cell counting kit-8 (CCK-8) (Dojindo Molecular Technologies, Shanghai, China) according to the manufacturer’s protocol. Briefly, rat primary hippocampal neuron were plated in a 96-well plate at a density of 1×10^3^/100μl per well. CCK-8 was added to each well either immediately after radiation or 1 h after THz radiation. Cells were further incubated at 37°C for 4 h and absorbance at 450 nm was measured.

### 2.4 Cellular apoptosis

Immediately after exposure to various THz radiation, the cellular apoptosis were analyzed by Annexin V staining (Beijing Biosea Biotechnology, Beijing, China). Briefly, cells were washed and resuspended in 100 μl binding buffer. Each sample was incubated with 5 μl Annexin V-APC at the room temperature for 10 min followed by incubation with 5 μl propidium iodide for 5 min. Then, 400μl PBS was added and the cellular apoptosis were analyzed by flow cytometer (FACSCalibur, Bio-Rad, USA).

### 2.5 Transmission electron microscopy

Cultured hippocampal neurons were harvested by centrifugation immediately after THz radiation. Cells (1×10^6^/ml) were fixed in a solution containing 2.5% glutaraldehyde for 2 h followed by another 2 h fixation with 1% osmium tetroxide. Specimens were dehydrated by ethanol and acetone and embedded with Epon812. Ultrathin sections (70nm) were made and stained with uranyl acetate and lead citrate. The cellular structure was observed by a transmission electron microscope (HITACHI H7650, Japan).

### 2.6 Quantification of amino acid neurotransmitter by HPLC

Immediately after THz radiation, 500μl cell medium was aspirated and centrifuged at 3000 rpm for 10min at 4°C. The supernatant was aliquoted and stored at 4°C. The concentrations of amino acid in the supernatant were measured by HPLC (Hewlett Packard, USA). The amounts were calculated based on standard solutions.

### 2.7 Detection of PSD-95 by immunofluorescence

Cells were fixed using 4% paraformaldehyde in PBS (pH 7.4) for 10 min at room temperature and washed 3 times with cold PBS, followed by permeabilization by 0.2% Triton X-100 for 10-15 min at room temperature. Cells were incubated with PBST-BSA buffer (TBS with 0.1% Tween-20 and 3% BSA) for 30min at 37°C, and then the antibody against PSD-95 (Sigma, USA) was added to the coverslip. After incubation at 4°C overnight, samples were washed 3 times with PBST, 5 min each wash. After incubated with diluted secondary antibody (1:200 in PBST) at room temperature for 45 min and 3 times wash with PBST, the coverslip was sealed with DAPI mounting medium (Sigma, USA). Slides were examined under Radiance 2100TM confocal microscope (Bio-Rad, USA).

### 2.8 Statistical analysis

All data were expressed as (mean±s.e.m). Statistical analysis was performed using SPSS18.0 software and two-tailed t-test. The difference with *p*<0.05 was considered as statistically significant.

## 3. Results

### 3.1 THz radiation alter the activity of primary hippocampal neuron depending on power density and exposure period

Hippocampus is critical for memory and learning, and the potential effects of THz radiation on rat hippocampal neurons were investigated in this study. The hippocampal neurons were identified by observing morphological characteristics and detecting MAP2 protein. MAP2 protein is a neuron-specific, microtubule-associated protein which could promote the assembly and stability of the microtubule network, and has been emerged as a marker of neuronal cells [8]. And, typical neuronal morphology could be detected by MAP2 staining (Fig 1A). The cellular activities were analyzed by cell counting kit-8 (CCK-8) assay in primary hippocampal neurons after exposure to various THz radiation. Compared to sham-radiated cells, the cellular activity was significantly decreased immediately and 1h after exposure to 0.12 THz with power intensity of 10 mW/cm^2^ for 10 min (Fig 1B). Similarly results were observed in primary cells after exposure to 0.12 THz with power intensity of 10 mW/cm^2^ for 30 min (Fig 1C). Moreover, the cellular activity of primary cells increased obviously after exposure to 0.157 THz with power intensity of 50 mW/cm^2^ for 30 min, but not for 10 min (Fig 1D-1E). Because exposure to 10 mW/cm^2^ and 50 mW/cm^2^ for 30 min exerted opposite effects on cellular activity, we investigate the effects of THz on primary hippocampal neurons by 30 min’s radiation.

**Figure 1.** Effect of THz radiation on activity of the primary hippocampal neuron. Primary rat hippocampal neurons were isolated and cultured as described in materials and methods. **(A)**, primary hippocampal cells showing neuronal staining with MAP2 (green). Nuclei were stained with DAPI (blue) (scale bar=50μm). Cellular activity was measured by CCK-8 assay immediately or 1 h after indicated THz radiation. The cellular activities were shown in B and C after exposure to 0.12 THz with power intensity of 10 mW/cm^2^ for 10 min and 30 min, respectively. Moreover, cellular activities were also presented in D and E after radiated by 0.157 THz with power intensity of 50 mW/cm^2^ for 10 min and 30 min respectively. Sham-radiated cells under the same conditions were used as the controls. Data were shown as mean±s.e.m. *,p<0.05, **,p<0.01 vs corresponding group.

### 3.2 THz radiation induced the cellular apoptosis in primary hippocampal neuron

The alteration of cellular activity might due to the changes of cell metabolism or cellular apoptosis. Therefore, we further analyzed the apoptosis by Annexin V and propidium iodide (PI) straining followed by flow cytometry. The results indicated that both 0.12 THz wave with power intensity of 10 mW/cm^2^ or 0.157 THz wave with power intensity of 50 mW/cm^2^ induced significant apoptosis (Fig 2 A-2B).

**Figure 2.** Effect of THz radiation on the apoptosis and ultrastruture of primary hippocampal neuron. Rat hippocampal neurons were radiated by 0.12 THz with power intensity of 10 mW/cm^2^ for 30 min or by 0.157 THz with power intensity of 50 mW/cm^2^ for 30 min. Apoptosis was analyzed by flow cytometry using Annexin V and propidium iodide staining immediately after THz radiation. The representative images were shown in **A**, and the statistical analysis were shown in B. Sham-irradiated cells were used as the controls. Data were shown as mean±s.e.m. **,p<0.01 vs sham group. The ultrastructure of the cell was examined by transmission electron microscopy (TEM) after exposure to sham-radiation (**C**), 0.12THz radiation with power intensity of 10 mW/cm^2^ for 30 min (**D**), or 0.157 THz radiation with power intensity of 50 mW/cm^2^ for 30 min (**E**). Organelles are indicated by arrows, green: nucleus; red: mitochondria; blue: ER; yellow: lysosome. Scale bars are 500nm in **C**, and 2µm in **D** and **E**.

Then, the ultrastructure was observed by transmission electron microscopy (TEM). Sham-radiated cells showed nuclei with homogenous chromatin, intact organelles and very few damaged mitochondrial cristae (Fig 2C). Cells treated with power intensity of 10 mW/cm^2^ (0.12 THz) showed nuclei with homogenous chromatin. However, swollen mitochondria, damaged cristae, swollen endoplasmic reticulum and increased lysosome could be detected (Fig 2D). Cells treated with power intensity of 50 mW/cm^2^ (0.157 THz) showed similar changes but with fewer lysosomes (Fig 2E). These results suggested that both 10 mW/cm^2^ and 50 mW/cm^2^ THz could cause induce structural injuries and result in cellular apoptosis.

### 3.3 THz radiation decreased the release of amino acid neurotransmitters in primary hippocampal neurons

Amino acid neurotransmitters are critical for transmitting nerve messages across the synapses. Some are inhibitory amino acids (IAAs) while others are excitatory amino acids (EAAs). The ratio of these amino acids could affect brain function[9]. In this study, we measured the concentrations of 16 amino acids in the sham-radiated and THz radiated primary hippocampal neurons by HPLC (Fig 3). In the culture medium, the concentration of Ala, Gly, Phe, His and Ser was not affected by THz radiation (Fig 3K-O), while the level of Val, Lue, Ile, Lys, Arg, and Thr was slightly down-regulated after exposure (Fig 3A-F). Moreover, the Met and Pro in 50 mW/cm^2^ radiated cells was lower than those in other two groups, and statistical difference were detected in comparing the concentration of Pro (Fig 3 G, H). Furthermore, THz radiation caused an increase in the concentration of Glu of Tyr although no statistical difference could be observed (Fig 3I-3J). Interestingly, Cyss level was significantly decreased after cells were radiated with 10 mW/cm^2^ of THz, but not in 50 mW/cm^2^ THz radiated cells (Fig 3P). Overall, THz radiation could slightly reduce the release of most amino acid neurotransmitters.

**Figure 3.** Effect of THz radiation on the release of amino acid neurotransmitters. Primary rat hippocampal neurons were radiated by THz wave at average power intensity of 10 mW/cm^2^ and 50 mW/cm^2^ for 30 min. The culture media were collected and the concentrations of 16 amino acids were analyzed by HPLC. Sham-radiated cells were used as the controls. Data were shown as mean±s.e.m. **,p<0.01 vs sham group.

### 3.4 THz radiation had no obvious effects on expression of PSD95 protein

To determine whether THz radiation would affect neuron function, we examined the expression of postsynaptic density protein (PSD)-95, an important scaffold protein promoting synapse maturation, affecting synaptic strength and plasticity[10]. We used immunofluorescence microscopy to observe the changes in the expression and localization of PSD-95. The expression pattern of PSD-95 in THz radiated (10 mW/cm^2^ or 50 mW/cm^2^) cells did not differ significantly from that in the sham-radiated cells (Fig 4).

**Figure 4.** Expression of PSD-95 protein in neurons after exposure to THz radiation in the cultured Primary rat hippocampal neurons were exposed to THz radiation at an average power intensity of 0, 10 or 50 mW/cm^2^ for 30 min. The expression and localization of PSD-95 (green) was examined by immunofluorescence microscopy.

## 4. Discussion

THz has been widely used in braining imaging, however its potential effects on neuronal cells are still largely unexplored. In models of shellfish neurons and mollusk neurons, it has been reported that THz laser radiation caused damage to membrane integrity and membrane potential respectively [11] [5]. In this study, we investigated the effects of THz radiation on morphology, cellular activity, apoptosis, and the release of neurotransmitter in primary neuronal cells *in vitro*. We found that the effects are closely associated with the THz parameters, including power intensity, duration and wavelength. Generally, both 10mW/cm^2^ and 50mW/cm^2^ THz radiation induced cellular apoptosis obviously in neurons companying with damaged mitochondria and increased lysosomes. However, low power intensity radiation (10mW/cm^2^) decreased cellular activity, while high power intensity radiation (50mW/cm^2^) increased cellular activity. The contrary effects between 10 mW/cm^2^ and 50 mW/cm^2^ THz exposure might due to the differential gene expression. It has been demonstrated that THz radiation different THz intensities caused opposite gene expression patterns in mouse mesenchymal stem cells [12]. However, the underlying mechanisms are still need to be clarified by using microarray, RNA-seq, ChIP-seq, and so on. Moreover, increased activity after 50mW/cm^2^ THz exposure might be attributed to the up-regulated calcium ions and enhanced enzyme activity [13].

In this study, we showed that THz radiation affected the release of amino acid neurotransmitter by hippocampal neurons *in vitro*. Amino acid neurotransmitters are able to transmit messages across the synapse, and can be grouped into excitatory amino acids (EAAs) which could stimulate neurons activities, such as Asp and Glu, and inhibitory amino acids (IAAs)which could inhibited firing, such as Gly and GABA[14]. The balance between EAAs and IAAs is important for maintaining brain functions. Our data suggested that THz radiation could slightly increase Glu, an pivotal EAA of cognition, motor coordination and emotions[15]. And, several amino acids including Val, Arg, Ile, Leu, Pro, and Thr were down-regulated after THz irradiation. These amino acids are involved in cognitive behavior such as learning and memory. For example, nitric oxide (NO) which is synthesized from L-Arg has been implicated in the learning process and in memory formation[16].Moreover, it has been reported that branched-chain amino acids (BCAAs), such as Leu, Ilu and Val, could slightly improve brain functions both in animal experiments and clinical application [17]. In addition, BCAAs are precursors of Glu and GABA, which are critical to maintain balance of brain activities. L-proline can activate excitatory glutamate and inhibitory glycine receptors in cultured rat neurons [18]. It has been reported that patients with hyperprolinemia showed neurological disorders, including spatial memory deficit [19]. D-serine is a co-agonist of the N-methyl D-aspartate (NMDA) receptor to regulate NMDA receptor transmission, synaptic plasticity and neurotoxicity[20]. Threonine is a precursor of inhibitory neurotransmitter glycine. L-threonine increased glycine concentration in the rat central nervous system, as well as produced antispastic effect in human [21] [22]. Overall, exposure to THz radiation could alter the release of amino acid neurotransmitters, which might cause negative effects, such as increased anxiety on mouse behaviors [4].

In addition to neurotransmitters, the balance between neuronal excitation and inhibition is also could be regulated by molecules that control the assembly of neurotransmitter receptors, signal transduction and synapse maturation. PSD-95, containing 3 PDZ domains, an SH3 domain and a guanylate kinase region, belongs to the membrane-associated guanylate kinase family [23, 24]. PSD-95 has an important role in regulating protein trafficking and ion channel clustering, particularly on glutamate receptor clustering and function. It binds to the NR2 subunit of NMDA-type glutamate receptors and modulates NMDA receptor clustering, activity and signaling [25]. PSD-95 could not only enhance AMPA recruitment and excitatory synaptic response [26], but also interact with other molecules to regulate synapse morphology, maturation and plasticity [27]. In this study, we found that THz radiation does not affect the expression or localization of PSD-95, indicating that THz radiation modulated synaptic functions through a PSD-95 independent mechanism.

## 5. Conclusions

The effect of THz radiation on rat hippocampal neurons *in vitro* are closely related with the parameters of radiation. Generally, THz induced apoptosis and altered cellular activity of primary hippocampal neurons, as well as regulated release of amino acid neurotransmitter, which in turn affected the homeostasis of neuronal excitability.

## Declarations

### Ethics approval and Consent to participate

All the experiments and corresponding procedures were approved by the Ethics Committee of Beijing Institute of Radiation Medicine.

### Consent for publication

All the authors approve it for publication.

### Availability of data and materials

All data generated or analysed during this study are included in this blished article

### Authors’ contributions

Bo Wang and Ruiyun Peng conceived the experiments. Li Zhao and Ruhan Yi designed the study, analyzed the data and wrote the paper. Shengzhi Tan conducted the microwave radiation. Hui Wang, Jing Zhang, Haoyu Wang performed the HPLC, TEM, and IF. Xinping Xu conducted the image analysis. Ji Dong and Binwei Yao contributed reagents/materials. All the authors reviewed the manuscript.

## Acknowledgements

Not applicable

## Authors’ information

Li Zhao, Ph.D, female. Beijing Institute of Radiation Medicine, No. 27 Taiping Road, Haidian District, Beijing 100850, PR China. Email.

Ruhan Yi, female, Beijing Institute of Radiation Medicine, No. 27 Taiping Road, Haidian District, Beijing 100850, PR China. Email.

Liu Sun, female, Beijing Institute of Radiation Medicine, No. 27 Taiping Road, Haidian District, Beijing 100850, PR China. Email.

Yunliang Chi, male. Beijing Institute of Radiation Medicine, No. 27 Taiping Road, Haidian District, Beijing 100850, PR China. Email.

Shengzhi Tan, Ph.D, male. Beijing Institute of Radiation Medicine, No. 27 Taiping Road, Haidian District, Beijing 100850, PR China. Email.

Ji Dong, female. Beijing Institute of Radiation Medicine, No. 27 Taiping Road, Haidian District, Beijing 100850, PR China. Email.

Hui Wang, Ph.D, female. Beijing Institute of Radiation Medicine, No. 27 Taiping Road, Haidian District, Beijing 100850, PR China. Email.

Jing Zhang, Ph.D, female. Beijing Institute of Radiation Medicine, No. 27 Taiping Road, Haidian District, Beijing 100850, PR China. Email.

Haoyu Wang, Ph.D, female. Beijing Institute of Radiation Medicine, No. 27 Taiping Road, Haidian District, Beijing 100850, PR China. Email.

Xinping Xu, female. Beijing Institute of Radiation Medicine, No. 27 Taiping Road, Haidian District, Beijing 100850, PR China. Email.

Binwei Yao, male. Beijing Institute of Radiation Medicine, No. 27 Taiping Road, Haidian District, Beijing 100850, PR China. Email.

Bo Wang, Ph.D, male. Central Laboratory, Hainan General Hospital, Hainan Affiliated Hospital of Hainan Medical University, No. 19, Xiuhua Road, Xiuying District, Haikou 570311, PR China. E-mail

Rui-Yun Peng, Ph.D, female, Beijing Institute of Radiation Medicine, No. 27 Taiping Road, Haidian District, Beijing 100850, PR China. E-mail

